# Mechanoreceptor Piezo1-Mediated Interleukin Expression in Conjunctival Epithelial Cells: Linking Mechanical Stress to Ocular Inflammation

**DOI:** 10.1101/2024.06.24.600298

**Authors:** Seiya Fukuoka, Naoki Adachi, Erika Ouchi, Hideshi Ikemoto, Takayuki Okumo, Hidetoshi Onda, Masataka Sunagawa

## Abstract

**Purpose:** Mechanical stress on the ocular surface, such as from eye-rubbing, has been reported to lead to inflammation and various ocular conditions. We hypothesized that the mechanosensitive Piezo1 channel in the conjunctival epithelium contributes to the inflammatory response at the ocular surface after receiving mechanical stimuli.

**Methods:** Human conjunctival epithelial cells (HConjECs) were treated with Yoda1, a Piezo1-specific agonist, and various allergens to measure cytokine expression levels using qRT-PCR and Western blot. Piezo1 activation-induced intracellular signaling pathways were also investigated. Mechanical stretching experiments were conducted to simulate Piezo1 activation in HConjECs. In *in vivo* studies, using immunohistochemistry, rats were administered Yoda1 eye drops to examine the inflammatory response in the conjunctiva and Piezo1-induced signaling activation.

**Results:** HConjECs expressed functional Piezo1 channel, and its activation significantly increased IL-6 and IL-8 expression through the p38 MAPK-CREB pathway. Piezo1-induced [Ca^2+^]_i_ elevation was crucial for the production of IL-6. Mechanical stretching mimicked these effects. *In vivo*, Yoda1 administration led to enhanced immunoreactivity of phospho-p38 MAPK and phospho-CREB and increased IL-6 in the rat conjunctival epithelium. Significant neutrophil infiltration was also observed after Piezo1 channel activation without affecting eosinophil numbers.

**Conclusion:** Mechanical stress-induced Piezo1 channel activation in conjunctival epithelial cells can cause ocular inflammation by upregulating pro-inflammatory cytokines via the p38 MAPK-CREB pathway and promoting neutrophil infiltration. These findings suggest that mechanical stimuli on ocular surface tissues are significant risk factors for ocular inflammation.

**Highlights:**

- Piezo1 channel activation increased IL-6 via p38 MAPK-CREB pathway in Human conjunctival epithelial cells (HConjECs).
- Piezo1 activation mimicked fungal extract but not pollen or dust mite.
- Mechanical stretching mimicked Piezo1 effects, boosting IL-6 production in HConjECs.
- Piezo1 activation elevated p-p38 MAPK, p-CREB, and IL-6 in the rat conjunctiva.
- Piezo1 activation leads to neutrophil infiltration in the rat conjunctival epithelium.

**Graphical abstract:** 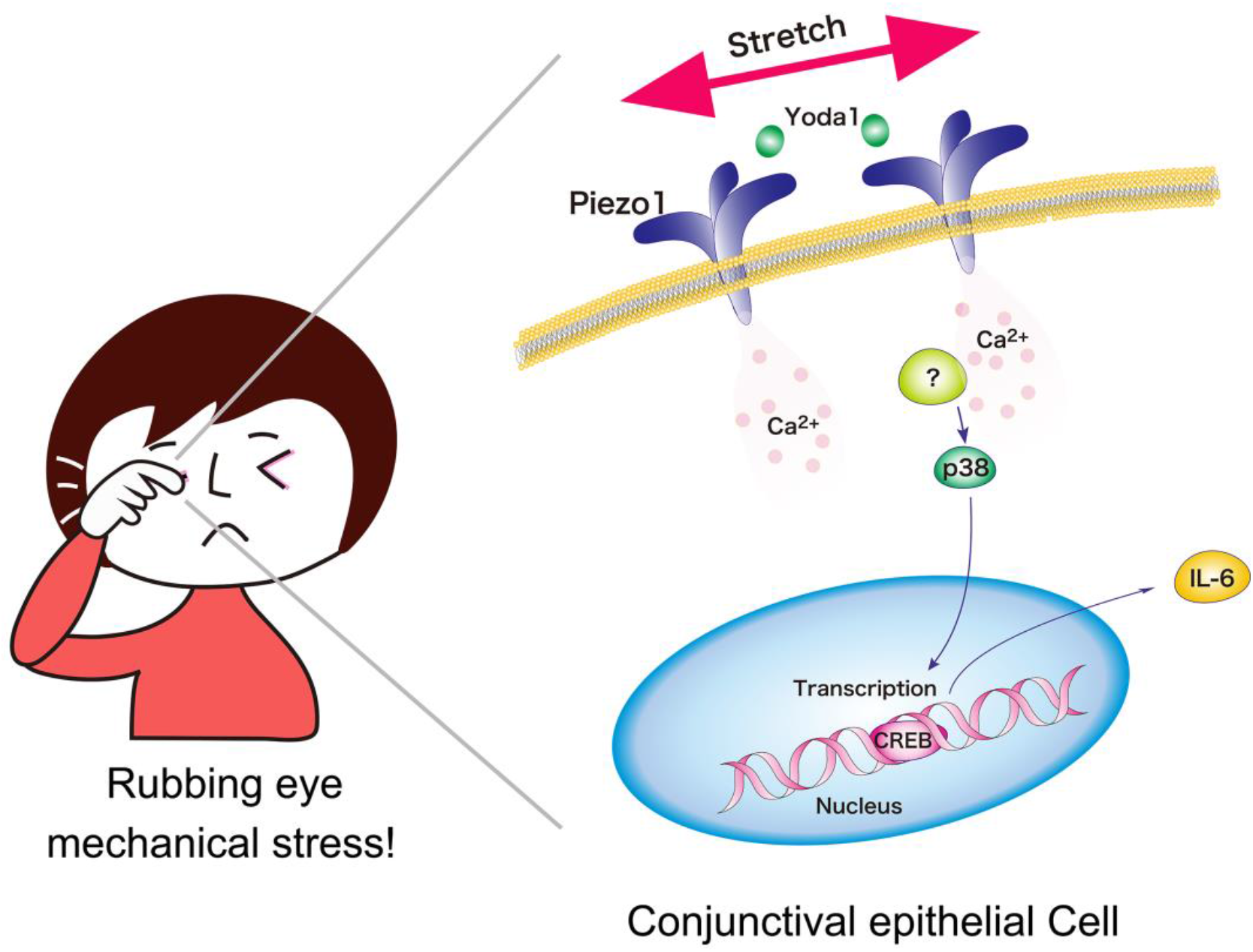

Mechanosensitive Piezo1 channel activation induces IL-6 expression in the conjunctival epithelial cells via elevation of [Ca^2+^]_i_ and activation of p38 MAPK and transcription factor CREB.

Mechanical stress stimulates Piezo1 channel and Ca^2+^ influx through it, which induces activation of p38 MAPK. A transcription factor CREB is phosphorylated after p38 MAPK activation, then promotes *IL-6* transcription.

## 1. Introduction

Rubbing eyes is a common reflex that many individuals engage in when experiencing discomfort or itchiness. Generally, individuals are advised not to rub their eyes to prevent infections from fingers and direct damage to the ocular surface which can lead to conditions such as cataracts and retinal detachment. However, recent studies have shown that eye-rubbing can significantly worsen various ocular conditions. As eye-rubbing increases intraocular pressure, chronic habit of abnormal rubbing may exacerbate glaucoma, leading to further optic nerve damage and irreversible vision loss [1]. Mechanical stress like eye-rubbing has been reported to cause corneal thinning, a condition known as keratoconus [1,2].

Recent clinical and meta-analysis studies have concluded that rubbing the eyes leads to the thinning of keratocytes, and the extent of this effect is influenced by the duration and intensity of the eye-rubbing [[3], [4], [5],[6]]. Interestingly, Mazharian *et al*. reported that, in addition to eye-rubbing, sleeping positions that involve pressing the eye against a pillow or sleeping surface were observed in patients with unilateral or highly asymmetric keratoconus [7]. Harrison *et al*. found that atopic keratoconus patients more often developed the disease on the side of their dominant hand [8]. Despite the clear involvement of eye rubbing in the onset and progression of keratoconus, its underlying mechanisms remain largely unknown. Many recent studies, however, strongly suggest that keratoconus is likely an inflammatory disease [[9], [10], [11]].

Ocular inflammation is a multifaceted response that is often triggered by various irritants. Common allergens and pathogens such as pollen, pet dander, fungi, dust mites, and viruses, and, as well as environmental stimuli like smoke and chemical substances, can provoke immune responses on the ocular surface. When the eyes are exposed to these agents, epithelial cells and resident immune cells release pro-inflammatory cytokines and chemokines. These cytokines and chemokines play a pivotal role in mediating the inflammatory response by promoting the recruitment and mutual activation of immune cells at the site of inflammation [12,13].

Most studies examining the tears of keratoconus patients and those wearing contact lenses—which are considered to induce mechanical stress on the ocular surface tissues—have identified elevated levels of pro-inflammatory cytokines such as interleukin-1β (IL-1β), interleukin-6 (IL-6), tumor necrosis factor-α (TNF-α), and matrix metalloproteinase-9 (MMP-9) [[14], [15], [16], [17]]. Ocular surface immune cells, such as neutrophils and natural killer (NK) cells also have been reported in the patients of keratoconus [18]. Importantly, Balasubramanian et al. have demonstrated that eye-rubbing itself acutely induced the release of cytokines such as IL-6, IL-8, and TNF-α, which play crucial roles in ocular surface inflammation, in the tears of normal subjects [19]. It is, therefore, possible that the mechanical stress caused by eye-rubbing and wearing contact lenses can induce inflammatory responses at the ocular surface through the activation of mechanosensitive ion channels.

The Piezo family of ion channels, comprising primarily Piezo1 and Piezo2, plays a crucial role in mechanotransduction—the process by which cells convert mechanical stimuli into electrochemical activity—and functions as cation channels that mainly permeate Ca^2+^, responding to physical changes such as pressure, stretch, and shear stress [20, 21]. Piezo1 is predominantly expressed in non-neuronal tissues, including blood vessels and internal organs, where it regulates physiological functions like vascular tone and red blood cell volume.Conversely, Piezo2 is mainly found in sensory neurons, where it mediates the sensations of touch, pain, and proprioception [22]. Interestingly, recent studies have started to report that Piezo1 channels are implicated in inflammatory responses and various pathological conditions by inducing cytokine production driven by mechanical stress, highlighting their roles beyond mechanosensation. [[23], [24], [25], [26], [27], [28], [29]]. However, the function of Piezo1 in ocular tissues and its relationship to ocular inflammation remains unclear.

This study aims to elucidate the previously suspected but not clearly understood phenomenon that mechanical stress can induce or exacerbate ocular inflammation, focusing on the role of the Piezo1 channel in conjunctival epithelial cells and its underlying mechanisms.

## Materials and Methods

### 2.1. Animals

Seven-week-old male Wistar rats were obtained from Nippon Bio-Supp. Center, located in Tokyo, Japan. The rats were kept in groups of two to three per cage under controlled conditions: a 12-hour light/dark cycle with lights on at 7 am, a temperature of 25 ± 1°C, and humidity maintained at 40-50%. They had unrestricted access to food (CLEA Japan, CE-2, Tokyo, Japan) and water for a period of 10 days prior to the commencement of the experiments.All procedures involving animals were reviewed and approved by the Committee of Animal Care and Welfare of Showa University (Animal Experiment Approval Number: 05096). The experiments were conducted following the guidelines set forth by the same committee. Each experiment was conducted more than twice, with careful measures taken to minimize animal distress and to reduce the number of animals utilized in the research.

### 2.2. Cells, Allergens, and Drugs

Human conjunctival epithelial cells (HConjECs; ScienCell Research Laboratories, San Diego, CA) were utilized for experiments. The cells were cultured in plastic 35 mm dishes coated with ε-Poly-L-lysine (Cosmo Bio Co., LTD, Tokyo, Japan) to enhance cell adhesion and growth. The culture medium (ScienCell Research Laboratories) was maintained at standard conditions, ensuring optimal cell proliferation and viability. Each *in vitro* experiment was performed with at least two independent cultures with two different lots of HConjECs.

For allergen exposure experiments, extracts of common allergens were sourced from ITEA (Tokyo, Japan). The allergens included cedar pollen, *Alternaria alternata* (a type of fungi), and house dust mite. These were prepared at a final concentration of 50 µg/ml.

Regarding the pharmacological treatments, Yoda1 (Selleck Chemicals, Houston, TX) was used to activate mechanosensitive Piezo1 channel.Additionally, to investigate the role of specific signaling pathways, HConjECs were pretreated with SB203580 (Selleck Chemicals), a specific inhibitor of p38 MAPK, at a concentration of 10 µM. This pretreatment was performed for 20 minutes prior to the addition of Yoda1. Similarly, cells were pretreated with BAPTA-AM (Dojindo Laboratories, Kumamoto, Japan) a calcium chelator, for 40-50 minutes before Yoda1 administration.

### 2.3. Western Blotting

Cultured cells were lysed in a lysis buffer composed of 1% sodium dodecyl sulfate (SDS), 20 mM Tris-HCl (pH 7.4), 5 mM ethylene-diamine-tetraacetic acid (pH 8.0), 10 mM sodium fluoride, 2 mM sodium orthovanadate, 1 mM phenylmethylsulfonyl fluoride, and 0.5 mM phenylarsine oxide. The homogenate underwent centrifugation at 15,000 rpm for 30 minutes at 25°C, with the supernatant subsequently collected. Protein concentration was standardized using a bicinchoninic acid protein assay kit (Thermo Fisher Scientific, MA, USA). Each sample (10 μg of protein) was separated by SDS-PAGE on 8 or 10% gels and then transferred to a polyvinylidene difluoride membrane.

The membrane was blocked with 5% (w/v) BSA (Fujifilm Wako Pure Chemical, Tokyo, Japan) for one hour at room temperature before being incubated overnight at 4°C with the following primary antibodies: anti-phospho-p38 (1:1,000, #9211, Cell Signaling Technology, Billerica, MA); anti-p38 (1:1,000, #9212, Cell Signaling Technology); anti-phospho-CREB (1:1,000, #9198, Cell Signaling Technology); anti-CREB (1:1,000, #9197, Cell Signaling Technology); anti-IL-6 (1:200, #GTX110527, Genetex Co, CA, USA); anti-Piezo1 (1:200, #15939-1-AP, Proteintech, IL, USA); anti-phospho-STAT3 (1:1,000, #9145, Cell Signaling Technology); anti-STAT3 (1:1,000, #9139, Cell Signaling Technology); anti-NF-kB p65 (1:1,000, #AB32536, abcam, MA, USA); anti-phospho-NF-kB p65 (1:1,000, #AB86299, abcam); anti-βactin (1:5,000, #A3854, Sigma-Aldrich Japan Co., Tokyo, Japan)

Post-incubation, the membrane was washed with Tris-buffered saline containing Tween 20 (Sigma-Aldrich Japan Co.) and incubated with either a goat anti-mouse secondary antibody conjugated with horseradish peroxidase (1:1000, Jackson ImmunoResearch, PA, USA) or a goat anti-rabbit secondary antibody (1:1000, Rockland Immunochemicals, Gilbertsville, PA, USA) for two hours at room temperature. Chemiluminescence detection was performed using Pierce ECL Western blotting substrate (Thermo Fisher Scientific), and the resulting images were captured with an Ez-Capture MG system (Atto Co., Tokyo, Japan). Band intensities were quantified using Lane & Spot Analyzer software (Atto Co.).

### 2.4. Human Piezo1 mRNA Expression Analysis

Total RNA was extracted from cultured HConjECs using the RNeasy Micro Kit (Qiagen), and mRNA expression was analyzed via reverse transcription polymerase chain reaction (RT-PCR). Briefly, total RNA was isolated, and cDNA was synthesized using ReverTraAce (TOYOBO, Osaka, Japan) with an oligo dT primer. The PCR amplification of human Piezo1 was performed using a Veriti Thermal Cycler (Thermo Fisher Scientific, Waltham, MA, USA) and ExTaq (Takara, Shiga, Japan). The cycling conditions for the PCR included 30 cycles at 94 °C for 30 seconds, 57 °C for 30 seconds, and 72 °C for 45 seconds.

The primer sequences for Human Piezo1 (KC602455.1) were as follows: forward primer 5′-TGCATCTACTTCGCCCTGCT -3′ and reverse primer 5′-ATGGTGAACAGCGGCTCATA -3′.

The PCR products were separated by agarose gel electrophoresis and visualized using SYBR Gold Nucleic Acid Gel Stain (Thermo Fisher Scientific). Images were captured with a WUV-M20 Printgraph Classic UV transilluminator (ATTO CORPORATION, Tokyo, Japan).

### 2.5. Cell Stretching Experiment

Stretching chambers (SC-0040; STREX Inc., Osaka, Japan) were pre-coated with 0.2% polyethyleneimine (Wako pure chemical, Tokyo, Japan) before seeding the cells. HConjECs were then seeded into these chambers and incubated for 24-48 hours in a humidified environment containing 5% CO_2_ at 37 °C until they reached confluence. Following the incubation period, the cells were subjected to a 120% stretch using a stretch device (STB-100-10; STREX Inc.) for 15 minutes. After the stretching protocol, cells were collected for analysis using quantitative real-time PCR (qRT-PCR) and western blotting to evaluate changes in gene and protein expression.

### 2.6. Quantitative Real-Time Polymerase Chain Reaction (qRT-PCR)

Total RNA was isolated from HConjECs samples utilizing the RNeasy Fibrous Tissue Mini Kit (Qiagen, Hilden, Germany). Following RNA extraction, reverse transcription was conducted using the ReverTra Ace RT Master Mix with gDNA remover (TOYOBO, Osaka, Japan). The resultant cDNA was combined with PowerTrack SYBR Green Master Mix (Applied Biosystems, Waltham, CA, USA) and specific gene primers. The qPCR assays were carried out on a QuantStudio 3 Real-time PCR system (Applied Biosystems).

Primers used in the study are follows. For human IL-6: forward primer 5′-AAGCCAGAGCTGTGCAGATGAGTA -3′ and reverse primer 5′-TGTCCTGCAGCCACTGGTTC - 3′; human IL-8: forward primer 5′-ACACTGCGCCAACACAGAAATTA -3′ and reverse primer 5′-TTTGCTTGAAGTTTCACTGGCATC -3′; human GAPDH: forward primer 5′-TGCACCACCAACTGCTTAGC -3′ and reverse primer 5′-GGCATGGACTGTGGTCATGAG -3′.

Data were normalized by Efficiency-corrected Relative Quantification ΔΔCt Method with QuantStudio Design & Analysis software (Applied Biosystems), using a series of concentrations of template DNA prepared through serial dilutions. Standard curves were constructed by plotting Ct values against the logarithm of the relative dilution factors. The slopes of these standard curves were used to estimate amplification efficiencies. The Ct values for IL-6, IL-8 and GAPDH were obtained for each sample, and ΔCt values were calculated by subtracting the Ct value of the reference gene (GAPDH) from that of the target gene. Efficiency-corrected ΔΔCt values were then calculated to compare gene expression levels between experimental and control groups. The relative expression levels of interleukins were determined based on these efficiency-corrected ΔΔCt values.This efficiency-corrected relative expression level was used for subsequent analysis.

### 2.7. Calcium Imaging Using Fluo-8 in Cultured HConjECs

Calcium imaging was performed on cultured HConjECs using the fluorescent calcium indicator Fluo-8 AM (abcam). Cells were seeded onto 35 mm glass-bottom dishes (Matsunami Glass, Osaka, Japan) coated with ε-Poly-L-lysine coating solution (Cosmo Bio co. LTD.). Prior to imaging, the cells were incubated with the Fluo-8 AM (final concentration of 5 µM) at 37°C for 30-45 minutes. Calcium imaging was performed using a fluorescence microscope (Axio Observer, Carl Zeiss, Oberkochen, Germany) equipped with appropriate filters for Fluo-8.

### 2.8. Immunohistochemistry and Immunocytochemistry

After Yoda1 (5 µM) eye drops, rats were deeply anesthetized at indicated time points using a combination of three anesthetics administered intraperitoneally: medetomidine hydrochloride (0.3 mg/kg, Domitol; Nippon Zenyaku Kogyo Co., Ltd., Fukushima, Japan), midazolam (4.0 mg/kg, Sandoz; Sandoz K.K. Tokyo, Japan), and butorphanol (5.0 mg/kg, Vetorphale; Meiji Seika Pharma Co., Ltd., Tokyo, Japan). The rats were then intracardially perfused first with cold phosphate-buffered saline (PBS) at pH 7.4, followed by 4% paraformaldehyde in 0.1 M PBS. The tissue was then equilibrated in 30% sucrose for two days and subsequently stored at −80°C. Using a cryostat (CM1860; Leica Biosystems, Germany), the tissue was sectioned into 20 μm thick slices for immunostaining.

For immunostaining, the sections were rinsed three times in PBS and incubated in a blocking solution containing 10% goat serum and 0.3% Triton X-100 in PBS. The sections were then incubated with primary antibodies: anti-phospho-p38 (1:500, #4511, Cell Signaling Technology); anti-phospho-CREB (1:1,000, #9198, Cell Signaling Technology); anti-IL-6 (1:500, #12912, Cell Signaling Technology); anti-Myeloperoxidase (1:500, # A039829, Dako, Denmark); anti-Eosinophil Peroxidase (1:500, # MAB1087-I, Sigma-Aldrich Japan Co.), for 48 hours. After three PBS washes, the sections were incubated with AlexaFluor 488-conjugated secondary antibodies (1:200, Invitrogen, CA, USA), followed by nuclear staining with DAPI (Sigma Chemical Co.). Immunofluorescent images were obtained with Olympus FV1000D confocal microscope (Olympus, Tokyo, Japan).

HConjECs were cultured on 35 mm glass-bottom dishes (Matsunami Glass, Osaka, Japan) coated with ε-Poly-L-lysine coating solution (Cosmo Bio co. LTD.). The cells were maintained in an environment of 5% CO_2_ and 95% air at 37 °C. For fixation, HConjECs were treated with 4% paraformaldehyde containing 4% sucrose (Sigma Chemical Co.) for 20 minutes. Post-fixation, the cells were permeabilized with 0.2% Triton X-100 in PBS (Sigma Chemical Co.) for 5 minutes and then blocked with 10% goat serum in PBS for 1 hour. Subsequently, the cells were incubated overnight at 4 °C with anti-phospho-p38 (1:500, #4511, Cell Signaling Technology), anti-phospho-CREB (1:1,000, #9198, Cell Signaling Technology), or anti-IL-6 (1:500, #12912, Cell Signaling Technology), following an isotype-specific secondary antibody conjugated with Alexa 488 (1:200, Molecular Probes, CA) binding. Nuclear staining with DAPI (Sigma Chemical Co.) was conducted after secondary antibody binding.Fluorescent images were captured using a fluorescent microscope (Axio Observer, Carl Zeiss).

### 2.9. Statistics

All values are presented as means with their standard errors (SEM). To assess the distribution of each data set, the Shapiro-Wilk normality test was initially employed. For data exhibiting normal distribution, one-way ANOVA followed by Tukey’s multiple comparisons test was conducted. In cases where the data did not follow a normal distribution, the Kruskal-Wallis test followed by the Steel-Dwass test was applied. Statistical analyses were performed using EZR software (Saitama Medical Center, Jichi Medical University, Saitama, Japan) [30]. A *p*-value of less than 0.05 was considered to indicate statistical significance.

## Results

### 3.1 Piezo1 Activation Mimicked Fungal Response by Inducing ILs, p38 MAPK, and CREB in HConjECs

We first examined whether activation of the mechanosensitive Piezo1 channel changes the expression of inflammatory cytokines. Yoda1, a specific agonist for Piezo1, and several allergens were applied to cultured HConjECs. Surprisingly, Yoda1 clearly increased *IL-6* and *IL-8* mRNA levels in HConjECs to levels similar to those induced by fungal extract (*Alternaria alternata*) **(Fig. 1A, B)**. It is notable that the extracts of cedar pollen and house dust mite caused only weak or no changes in the expression of these interleukins **(Fig. 1A, B)**. Next, we investigated the intracellular signaling proteins activated after Yoda1 treatment. We found that p38 MAPK and the transcription factor CREB were activated (phosphorylated) **(Fig. 1C)**, while no activation of other representative signaling proteins involved in inflammatory responses, such as NF-κB and STAT3, was observed (data not shown). Among the agents that induced interleukin expression, both Yoda1 and fungal extract activated p38 MAPK and CREB, but Yoda1 appeared to have a stronger effect **(Fig. 1D)**. These data suggest that activation of the mechanosensitive Piezo1 channel can induce inflammatory cytokines, such as IL-6 and IL-8, in HConjECs at levels similar to those induced by fungal extract through common intracellular signaling pathways.

**Fig. 1.**
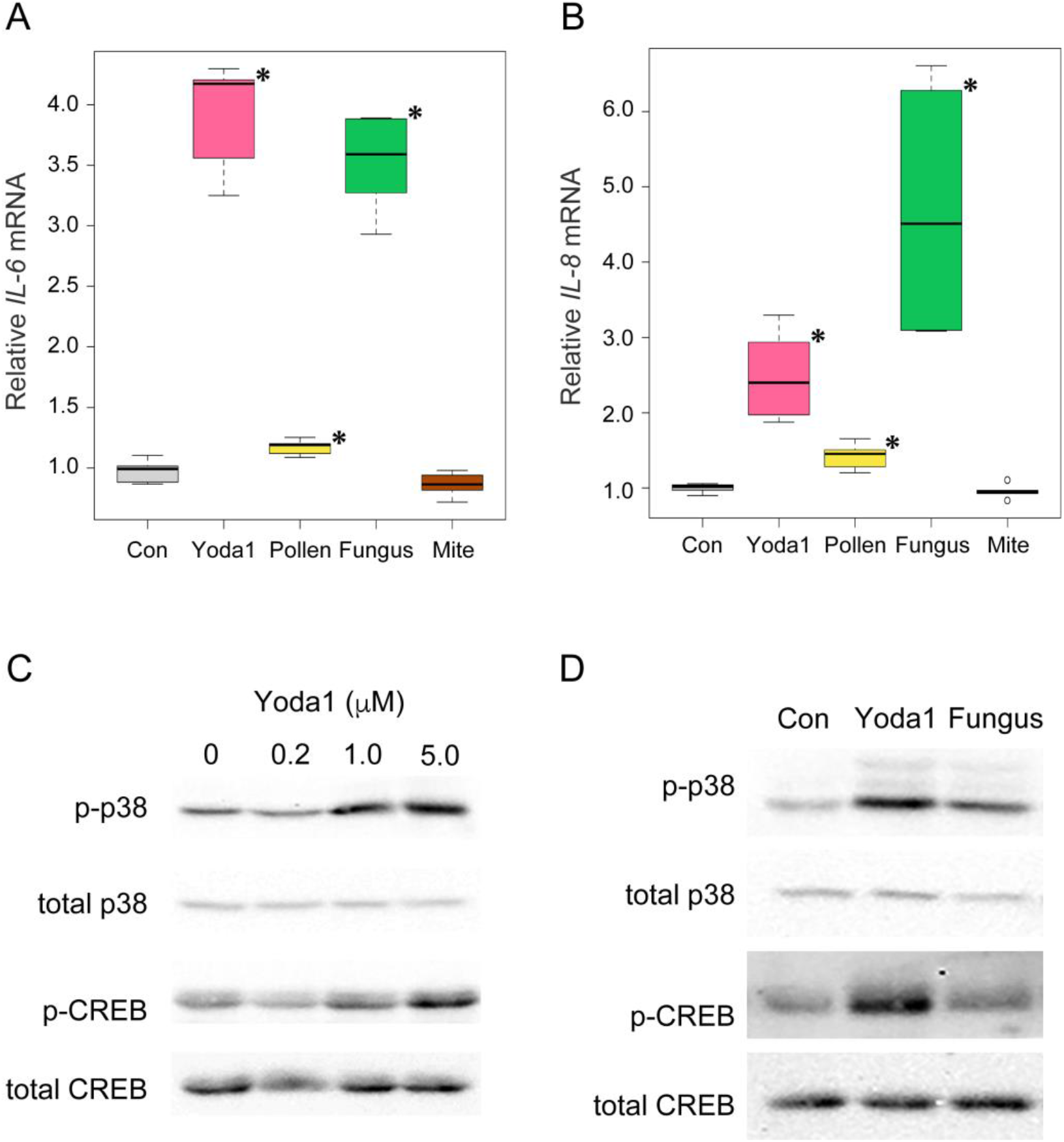
Piezo1 activation and Fungi extract induced IL-6 and IL-8 mRNA expression through p38 MAPK and CREB in HConjECs. HConjECs were treated by indicated allergens and piezo1 agonist Yoda1 for 6 hours. Relative mRNA expression levels of (A) IL-6 and (B) IL-8. Yoda1: 10 µM Yoda1; Pollen: 50 µg/ml extract of cedar pollen; Fungi: 50 µg/ml extract of Alternaria alternata; Mite: 50 µg/ml extract of house dust mite. (n=6) * *p* < 0.05 vs. Control group. (C) Dose-dependent activation of p38 MAPK and CREB by Yoda1. Yoda1 was treated for 15 min. (D) Both Yoda1 (5 µM) and Alternaria alternata extract activated p38 MAPK and CREB.

### 3.2 HConjECs Express Functional Mechanosensitive Piezo1 Channel

We next determined whether a functional Piezo1 channel was expressed in HConjECs. RT-PCR and western blot analysis clearly showed that HConjECs expressed the Piezo1 channel **(Fig. 2A, B)**. Imaging of intracellular Ca^2+^ demonstrated that [Ca^2+^]_i_ significantly increased immediately after stimulation of the Piezo1 channel by Yoda1 **(Fig. 2C, D)**. The elevation of [Ca^2+^]_i_ induced by Yoda1 lasted for at least 1 minute **(Fig. 2C, D)** (Supplementary Video 1). These data clearly indicate that HConjECs express a functional mechanosensitive Piezo1 channel.

**Fig. 2.**
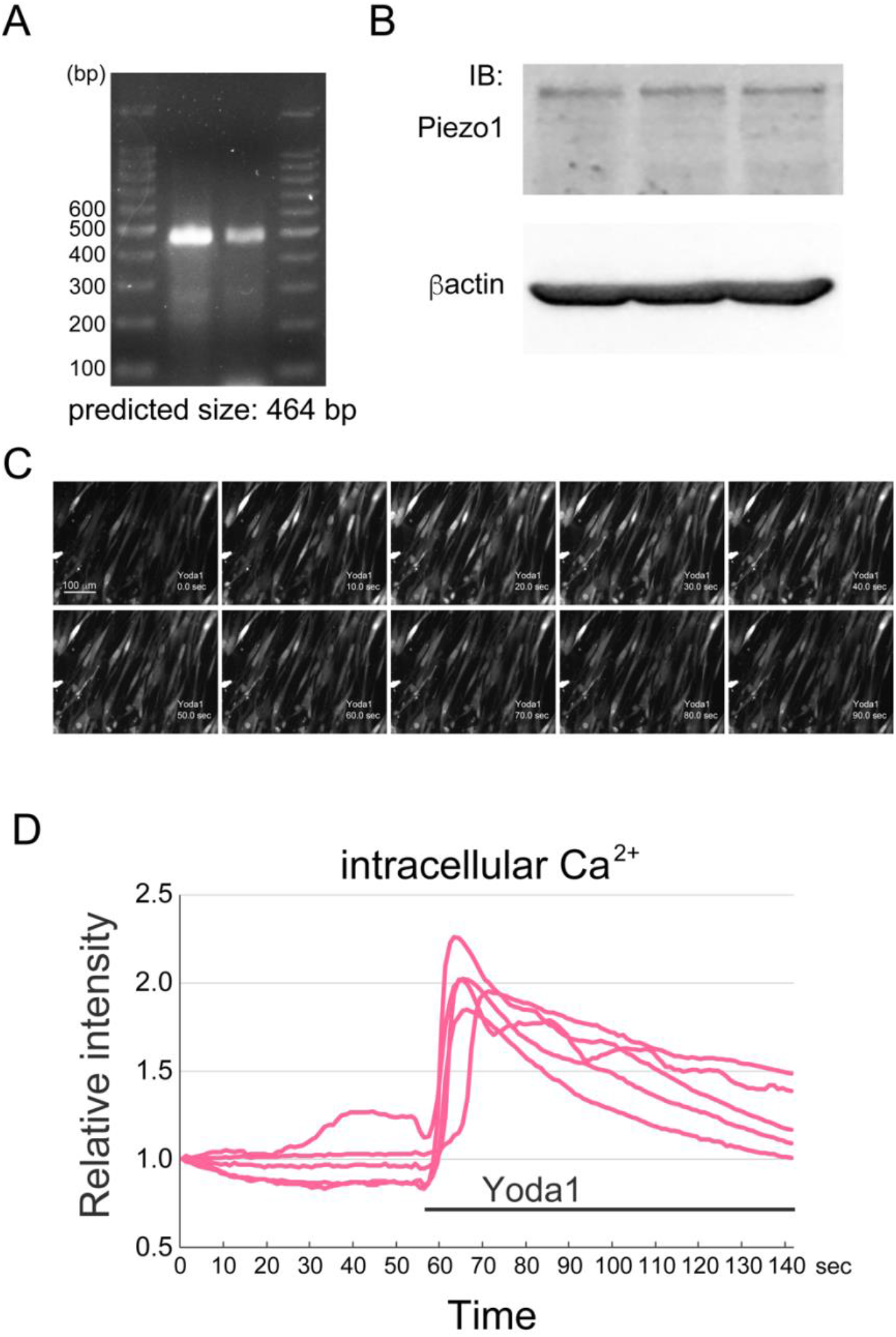
HConjECs express functional Piezo1. (A) RT-PCR with primers for a part of human PIEZO1 showed Piezo1 mRNA expression at the predicted size (464 bp). RNA samples of two independent culture from two different lots of HConjECs. (B) Piezo1 protein expression in HConjECs (predicted size of human Piezo1≒290 kD). Protein samples of three independent culture obtained from two different lots of HConjECs. (C) Yoda1 induced Ca^2+^ influx in HConjECs. Every 10 seconds image of Fluo-8 loaded HConjECs before and after Yoda1 exposure. (D) Time course of the relative intensity of [Ca^2+^]_i_ of five cells during Yoda1 exposure.

### 3.3 Piezo1-Induced IL-6 Upregulation via p38 MAPK-CREB Pathway and [Ca^2+^]_i_ Elevation

To confirm that Ca^2+^ influx through the Piezo1 channel and activation of p38 MAPK and CREB were involved in IL-6 upregulation, a specific inhibitor for p38 MAPK and a Ca^2+^ chelator were used. Pretreatment with SB203580, a specific inhibitor for p38 MAPK, successfully blocked the activation of p38 MAPK in HConjECs **(Fig. 3A, B)**. Importantly, both CREB activation **(Fig. 3A, C)** and IL-6 protein/mRNA upregulation **(Fig. 3D, E)** induced by Piezo1 channel activation were significantly suppressed by blocking p38 MAPK activation. Furthermore, chelating intracellular Ca^2+^ with BAPTA-AM significantly reduced Yoda1-induced IL-6 mRNA upregulation **(Fig. 3F)**. These data suggest that activation of the mechanosensitive Piezo1 channel induces IL-6 expression via Ca^2+^ influx and activation of the p38 MAPK-CREB pathway.

### 3.4 Nuclear p-p38 MAPK/p-CREB and Cytosolic IL-6 Induced by Piezo1 Activation

Immunocytochemical analysis showed that p38 MAPK and CREB were dramatically activated (phosphorylated) 15 minutes after Yoda1 treatment, and were found in the nuclei of HConjECs **(Fig. 4A, B)**. A surge of IL-6 protein-expressing cells was also observed 6 hours after Piezo channel activation **(Fig. 4C)**. Immunoreactivity of IL-6 was well co-localized with the *cis*-Golgi marker GM130 and showed a dot-like pattern outside of the *cis*-Golgi marker **(Fig. 4D)**. The activation of p38 MAPK and CREB, as well as the expression of IL-6 protein after Piezo1 activation, were clearly diminished by pretreatment with SB203580, a specific inhibitor for p38 MAPK **(Fig. 4A, B, C)**. These data indicate that activation of the Piezo1 channel induces phosphorylation and translocation of p38 MAPK to the nuclei, subsequently stimulating CREB activation and IL-6 upregulation.

**Fig. 3.**
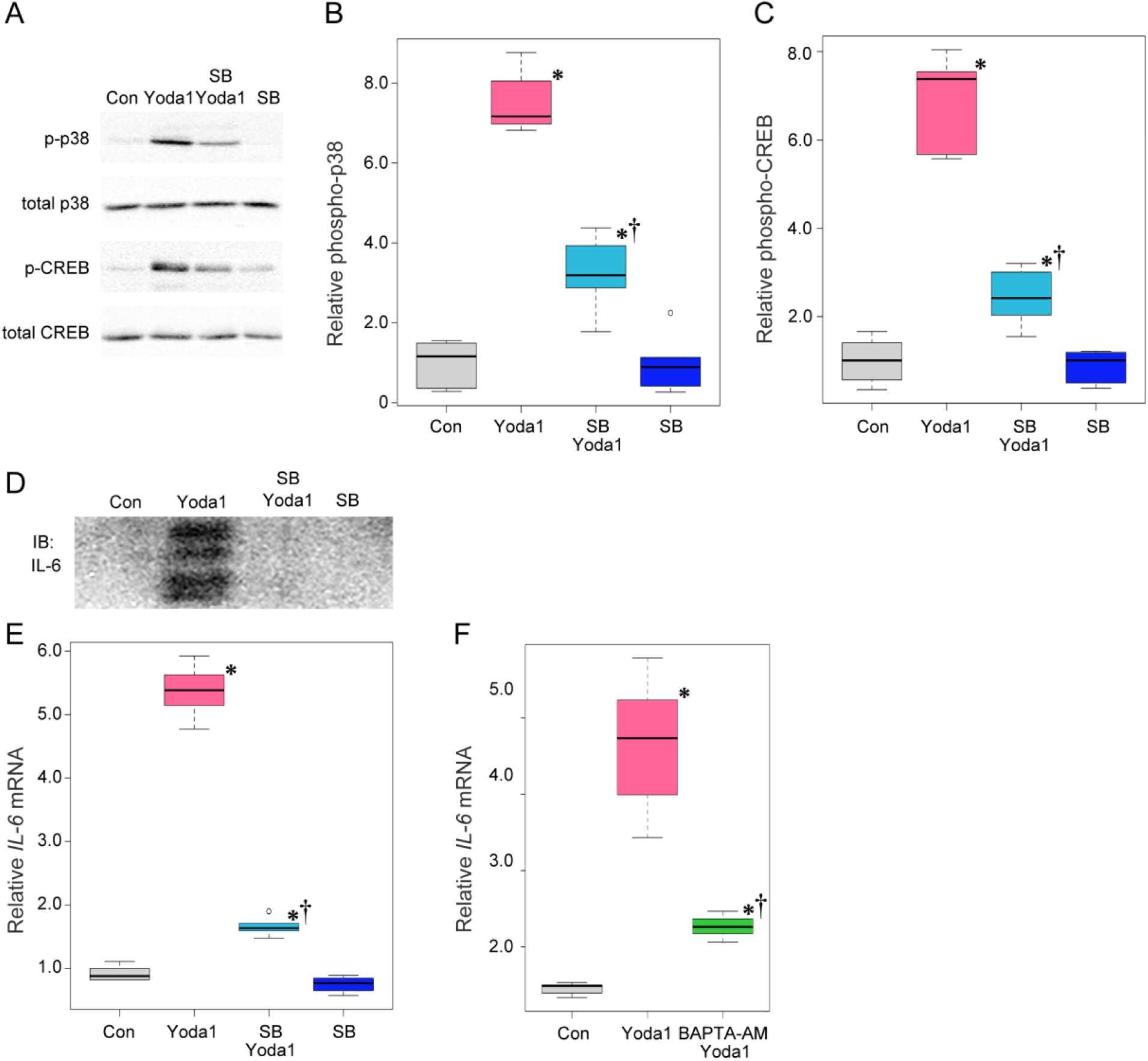
Activation of p38 MAPK and elevation of [Ca^2+^]_i_ were essential for Yoda1-induced IL-6 upregulation. (A, B, C) SB203580 (10 µM), a specific inhibitor for p38 MAPK, significantly suppressed activation of p38 MAPK (A, B) and CREB (A, C) after Yoda1 treatment. (D, E) Yoda1-induced IL-6 protein (D) and mRNA (E) upregulation (6 h) was completely dependent on p38 MAPK activation. (F) Chelating intracellular Ca^2+^ by BAPTA-AM blocked Yoda1-induced IL-6 mRNA upregulation. (n=6) *** *p* < 0.05 vs. Control group, †*p* < 0.05 vs. Yoda1 group.

**Fig. 4.**
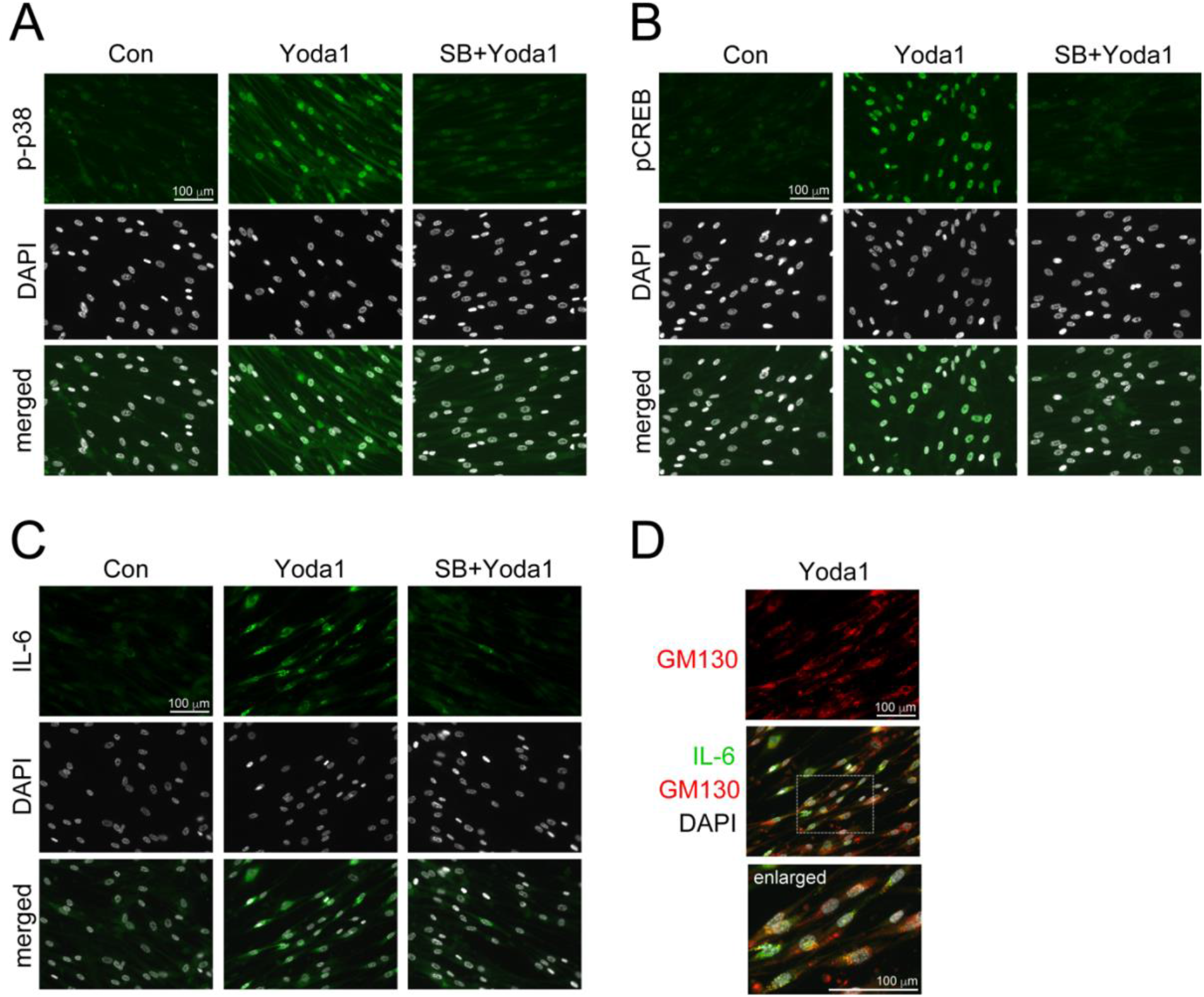
Piezo1 activation dramatically induced nuclear expression of phospho-p38 MAPK and phospho-CREB and cytosolic expression of IL-6 protein in HConjECs, which were critically dependent on p38 MAPK activation. (A) Yoda1-induced phosphorylation p38 MAPK and its nuclear expression (15 min after Yoda1 application). (B) Yoda1-induced phosphorylation CREB in nuclei (15 min). (C) Cytosolic IL-6 protein upregulation was induced by Yoda1 (6 h). (D) Co-localization of Yoda1-induced IL-6 protein and cis-Golgi marker GM130. Lower image was shown by enlarging the white dot square in middle merged image. SB: SB203580 (10 µM) was pretreated for 20 min. Scale bar = 100 µm.

### 3.5 Mechanical Stimulation Induced p38 MAPK/CREB Activation and IL-6 Upregulation in HConjECs

We next determined whether mechanical stimulation actually caused the phenomena induced by Yoda1-dependent Piezo1 channel activation. Cultured HConjECs were stretched to 120% and released 10 times per minute for 15 minutes **(Fig. 5A)**. The stretch stimulation induced activation of the p38 MAPK-CREB pathway without Yoda1 treatment in an extracellular Ca^2+^-dependent manner **(Fig. 5B, C)**. IL-6 upregulation was also confirmed 6 hours after mechanical stimulation in HConjECs, which was blocked by SB203580, a specific inhibitor for p38 MAPK **(Fig. 5D)**. These data show that Yoda1 treatment induces similar cellular responses as actual mechanical stress.

**Fig. 5.**
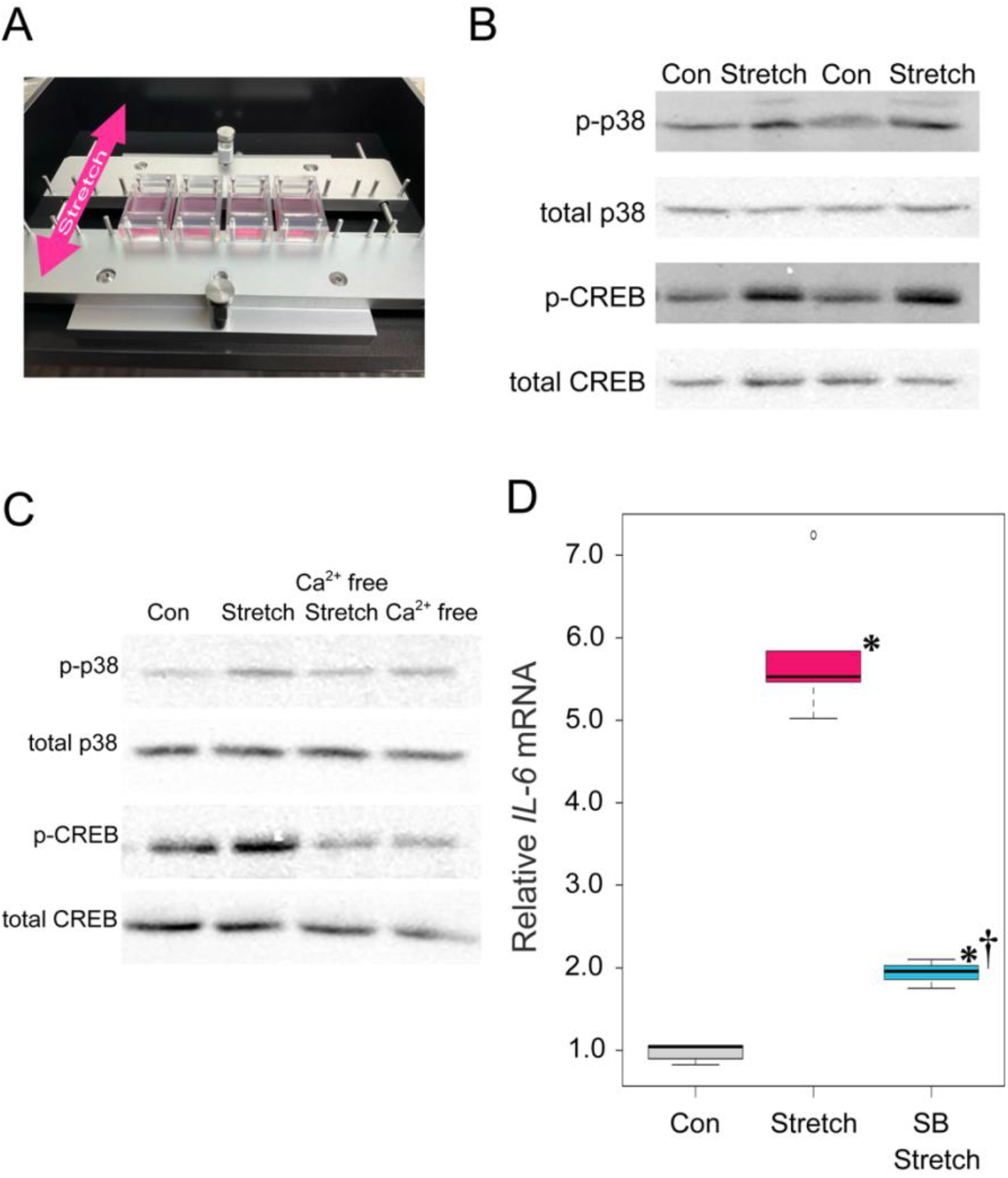
Mechanical stimulation induced p38 MAPK/CREB activation and IL-6 upregulation in HConjECs. (A) Cultured HConjECs were mechanically stimulated by 120% stretch and release cycles, performed 10 times per minute for 15 minutes. (B) Mechanical stimulation induced activation of p38 MAPK and CREB immediate after 15 minutes stimulation. (C) Activation of p38 MAPK and CREB were blocked by removing extracellular Ca^2+^ (Ca^2+^ free) 5 minutes before stretch stimulation. (D) Mechanical stimulation induced IL-6 mRNA (6 h) which was dependent on p38 MAPK activation. SB: SB203580 (10 µM) was pretreated for 20 min (n=6) * *p* < 0.05 vs. Control group, †*p* < 0.05 vs. Yoda1 group.

### 3.6 Piezo1 Activation Upregulates p-p38 MAPK, p-CREB, and IL-6 in The Rat Conjunctival Epithelium

Finally, we determined whether activation of the Piezo1 receptor in the rat conjunctival epithelium *in vivo* showed similar phenomena as observed in cultured HConjECs *in vitro*. A significant increase in expression of phospho-p38 and phospho-CREB in the rat conjunctival epithelium was observed 5-10 minutes after instillation of Yoda1 into the eye **(Fig. 6A, B, D, E)**. IL-6 immunoreactivity also dramatically increased 3 hours after Piezo1 channel activation **(Fig. 6C, F)**. These data confirmed that the activation of Piezo1 receptors in the rat conjunctival epithelium *in vivo* exhibited phenomena comparable to those seen in cultured HConjECs *in vitro*.

**Fig. 6.**
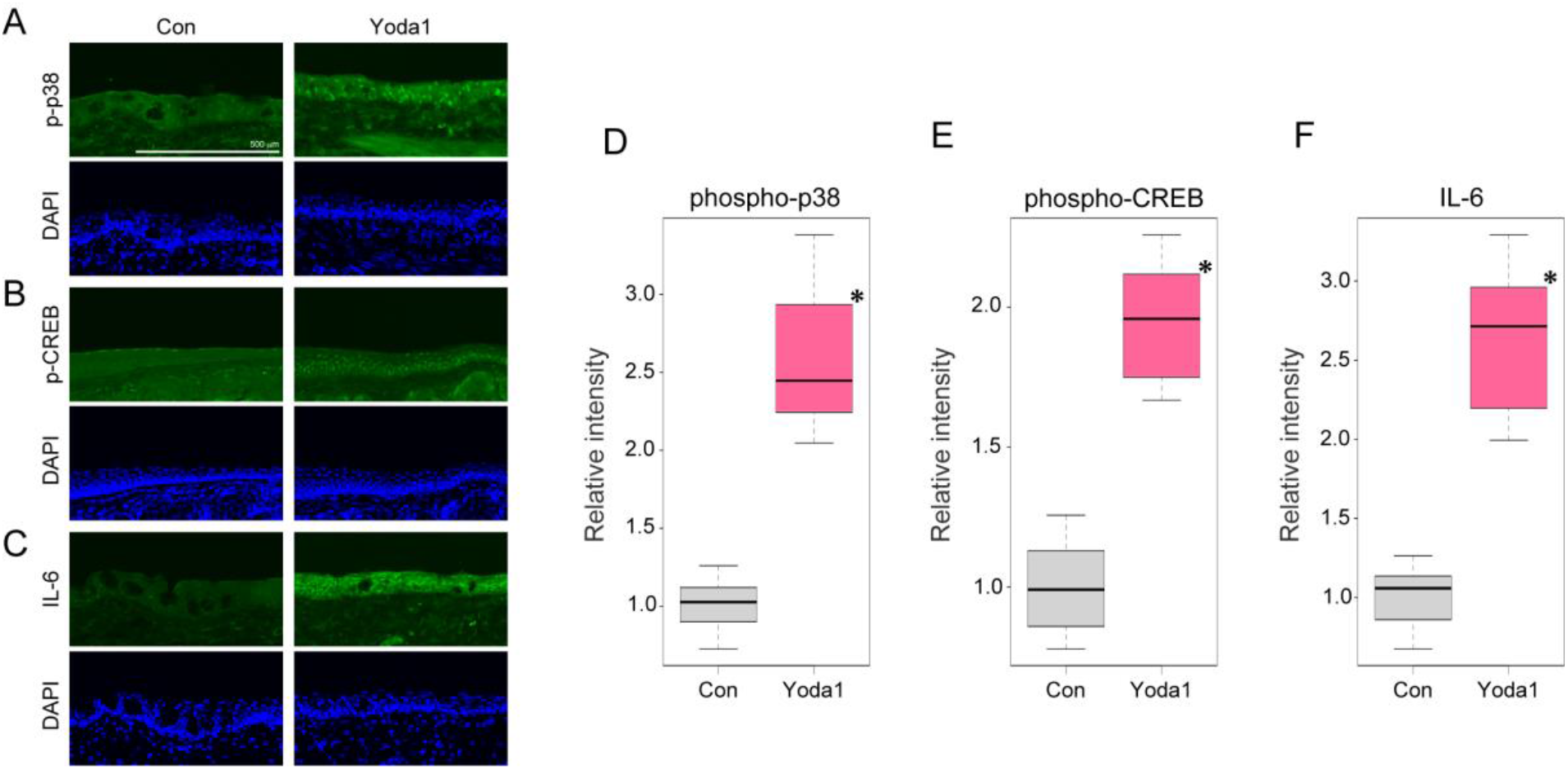
Piezo1 activation induced nuclear expression of phospho-p38 MAPK and phospho-CREB and upregulation of IL-6 protein in the rat conjunctival epithelium. Each rat was received instillation of Yoda1 to their one eye, then eye samples were collected 5-10 minutes, 3 hours after the instillation. (A-F) The instillation of Yoda1 (5 μM) to the eye induced nuclear expression of phospho-p38 MAPK (5-10 min, A, D) and phospho-CREB (5-10 min, B, E) and expression of IL-6 protein (3 h, C, F) in the rat conjunctival epithelium. (n=20) * *p* < 0.05 vs. Control group, †*p* < 0.05 vs. Yoda1 group.

### 3.7 Piezo1 Activation Promoted Neutrophil Infiltration into The Rat Conjunctival Epithelium

We also observed a significantly increased number of neutrophils, myeloperoxidase (MPO)-positive cells, 24 hours after Piezo1 receptor stimulation in the rat conjunctival epithelium **(Fig. 7A, C)**. However, the number of eosinophils, major basic protein (MBP)-positive cells, did not change 3 and 24 hours after instillation of Yoda1 into the eye (3 hours: data not shown; 24 hours: **Fig. 7B, D**). These data suggest that activation of the Piezo1 channel in the conjunctiva induces neutrophil infiltration into the conjunctival epithelium without affecting eosinophil infiltration.

**Fig. 7.**
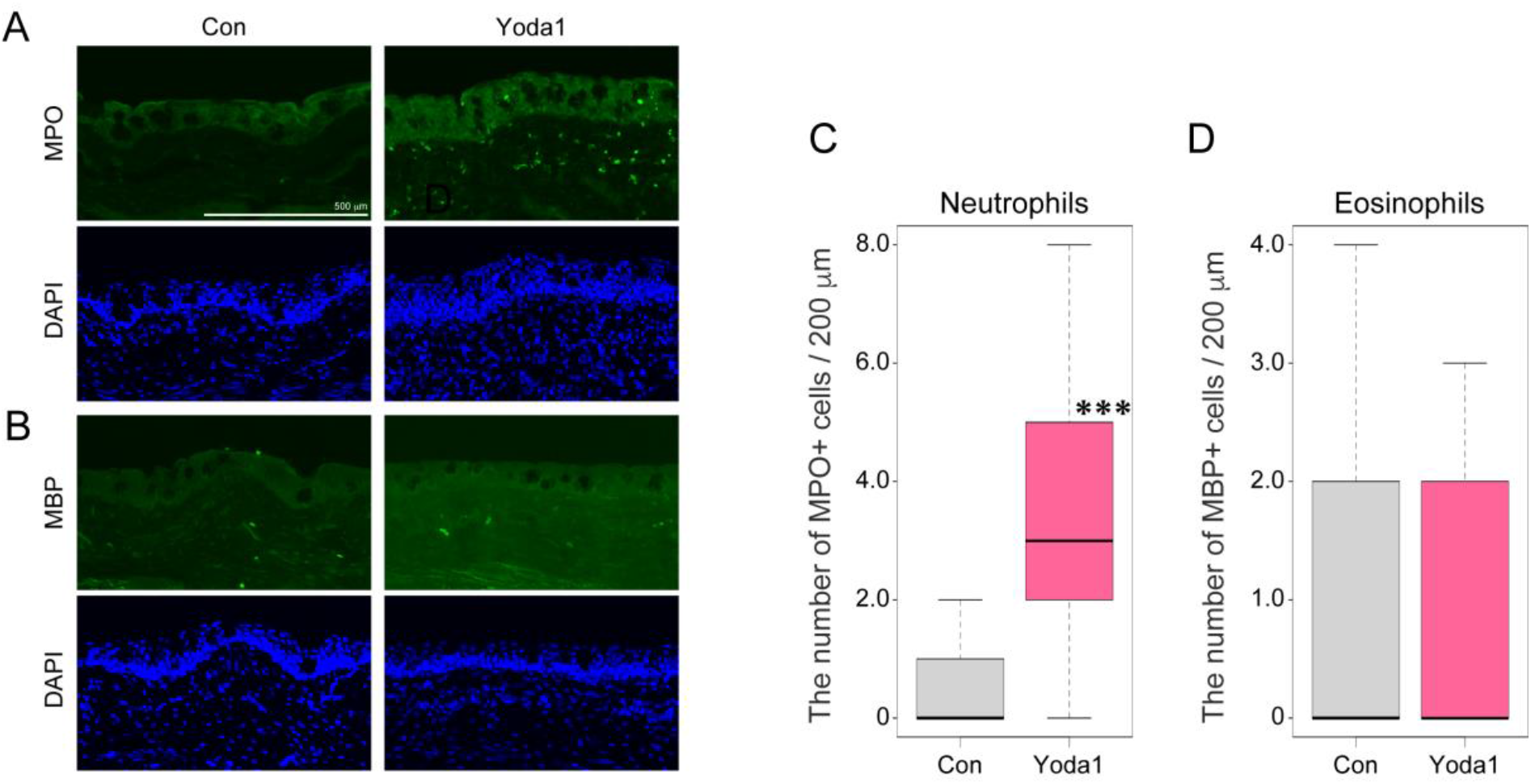
Piezo1 activation promotes neutrophil infiltration into the conjunctival epithelium in the rat eye. The eye samples were collected 24 hours after the instillation. (A, B) Sections of the rat conjunctiva were immunostained with markers for (A) neutrophils (MPO: Myeloperoxidase) and (B) eosinophils (MBP: Major Basic Protein). Intensity of immunoreactivity was measured in randomly selected regions of the conjunctival epithelium. (C, D) MPO-positive neutrophils (C) and MBP-positive eosinophils (D) were counted and quantified in randomly selected 200 µm parts of the conjunctival tissue. (n=20) * *p* < 0.05 vs. Control group, †*p* < 0.05 vs. Yoda1 group.

## 4 Discussion

In this study, it was demonstrated that mechanical stress could induce upregulation of pro-inflammatory cytokines involving IL-6 through Piezo1 mechanoreceptor channel-dependent Ca^2+^ influx and subsequent activation of the p38 MAPK pathway and a transcription factor CREB in conjunctival epithelial cells.

Upregulation of pro-inflammatory cytokines were strongly elicited by treatment with Yoda1 or a fungal extract but were either weak or absent with extracts of cedar pollen or house dust mite. It is remarkable that the activation of the Piezo1 channel induced an upregulation of *IL-6* mRNA to levels comparable to those caused by the extract of *Alternaria alternata*. Interestingly, both Yoda1 and extract of *Alternaria alternata* apeared to induce p38 MAPK and CREB activation. The results of these experiments using various allergens and infectious agents suggest that mechanical stress on the conjunctival epithelium could provoke inflammatory responses resembling infection rather than an allergic reaction.

Piezo1 channel has been found to be present in various tissues such as the lungs, bladder, and colon in vertebrates. Conversely, Piezo2 channel is primarily expressed in bone, sensory neurons, and dorsal root ganglion (DRG) neurons [31]. The expression of Piezo channels in ocular tissues has been documented in rodents, showing the presence of Piezo1/2 channels in the cornea and lens epithelial cells [32]. However, their expression in human ocular surface cells has not been clearly documented, which makes our findings significant: our results clearly showed that human conjunctival epithelial cells express functional Piezo1 channel.

The Piezo1 channel has been implicated in chronic inflammatory conditions associated with mechanical stress, activating various signaling pathways and leading to cytokine production [[23], [24], [25], [26], [27], [28], [29]]. In the circulatory system, Piezo1 channels are pivotal in cardiovascular development, blood pressure regulation, and vascular remodeling associated with hypertension. Piezo1 also plays a vital role in cardiovascular inflammation by detecting mechanical stress, leading to Ca^2+^ influx and the upregulation of IL-6 mRNA in cardiac fibroblasts [24]. Interestingly, this IL-6 mRNA upregulation in cardiac fibroblasts was p38 MAPK-dependent, consistent with our findings. Elevated IL-6 levels can enhance cardiac fibroblast proliferation and fibrosis, subsequently reducing myocardial compliance and pumping function [23].

In the respiratory system, Piezo1 activation by periodic hydrostatic pressure in the lungs induces Ca^2+^ influx and cJun phosphorylation in monocytes, leading to endothelin 1 (ET1) expression and subsequent activation of hypoxia-inducible factor 1α (HIF1α) via the endothelin receptor type B (EDNRB), promoting a prolonged pro-inflammatory response [25]. This process also increases chemokine CXCL2 secretion facilitating neutrophil migration and bacterial clearance in the lungs, which indicates essential role of Piezo1 for innate immunity [26]. On the other hand, *PIEZO1* gene knockout in mice demonstrated protective effects against bleomycin-induced pulmonary fibrosis [26].

Inflammatory responses involving the Piezo1 channel have also been reported in the skeletal system. Although Piezo1 channel has critical roles in bone development and mechano-stimulated bone homeostasis [27], Piezo1 significantly impacted inflammation by upregulating NLRP3 (nucleotide-binding domain, leucine-rich–containing family, pyrin domain–containing-3) inflammasome in cardiovascular disorders inflammasome, catabolic enzymes (MMP-9, MMP-13), and pro-inflammatory cytokine IL-1β, contributing to intervertebral disc degeneration (IDD) following mechanical stress in rat [28]. Piezo1 inhibition reduces NLRP3 and IL-1β activation in nucleus pulposus cells, mitigating mechanical shock-induced damage. Furthermore, in osteoarthritis, mechanical stress-mediated apoptosis of chondrocytes was attenuated via suppression of Piezo1 [29]. Activation of Piezo1 in nucleus pulposus cells increased Ca^2+^ load, triggering the NF-κB pathway, oxidative stress, and inflammation that can lead to lumbar degeneration [29].

Taken together, Piezo1 mechanoreceptor channel plays a crucial role in various organs by mediating Ca^2+^ signaling, which is essential for normal tissue development and pro-inflammatory responses necessary for maintaining innate immunity. However, excessive or prolonged activation of Piezo1 can lead to overactive immune response or apoptosis, thereby contributing to the development of immune-related diseases. Despite this, there have been few reports on the role of Piezo1 in epithelial tissues, including the eye and its surrounding structures. Current study represents the first evidence suggesting that Piezo1 activation in epithelial tissues promotes inflammatory responses.

As described above, downstream mechanisms of Piezo1exhibit notable variations in different tissue inflammations. Piezo1-mediated Ca^2+^ signaling influences the onset and progression of inflammation through multiple signaling pathways. Current study showed immediate activation of p38 MAPK and CREB after Piezo1-mediated Ca^2+^ influx to induce IL-6 expression in HConjECs.

The p38 MAPK is a crucial signaling molecule activated by various pro-inflammatory and stressful stimuli. Initially identified as a 38-kDa protein rapidly phosphorylated in response to lipopolysaccharide (LPS), a component of the outer membrane of Gram-negative bacteria, stimulation, p38 MAPK has been shown to induce cytokines such as IL-1 and TNF-α in LPS-stimulated monocytes.Due to its well-established role in inflammation, p38 MAPK is thought to serve as a potential therapeutic target in inflammatory diseases [33, 34]. The most extensively studied upstream activators of p38 MAPK are MAP kinase kinases (MKKs) such as MKK3 and MKK6. However, it is also known that p38 MAPK can be activated downstream of Ca^2+^ influx. Takeda *et al*. reported the involvement of apoptosis signal-regulating kinase 1 (ASK1), which is activated during TNF-α and endoplasmic reticulum (ER) stress, in the Ca^2+^-Ca^2+^/calmodulin-dependent protein kinase type II (CaMKII)-ASK1–p38 axis in fibroblasts [35]. Ye *et a*l. demonstrated the involvement of protein kinase C (PKC) in the phosphorylation of p38 MAPK downstream of Ca^2+^ influx in astrocytoma cells [36]. Although further detailed studies are necessary to elucidate the upstream factors of p38 MAPK activation in Piezo1-mediated Ca^2+^ signaling in HConjECs, these kinases might play a role in the signaling pathway. Although the association between p38 MAPK and ocular diseases has not been extensively reported, in an experimental dry eye model mouse, significant activation of p38 MAPK along with the production of IL-1β, TNF-α, and MMP-9 has been documented in the conjunctival and corneal epithelia [37].Interestingly, *in vitro* studies of dry eye model human corneal epithelial cells have shown not only p38 MAPK activation but also a notable increase in the expression of IL-6 and TNF-α [38].

While numerous transcription factors activated downstream of p38 MAPK have been reported [33], our study specifically found that p38 MAPK activated after Piezo1-dependent Ca^2+^ influx promotes *IL-6* transcription through CREB which can also be activated by other MAPKs and CaMKII [39]. In our experiments, inhibition of p38 MAPK by SB203580 strongly but not completely suppressed CREB phosphorylation or IL-6 expression, suggesting that, in HConjECs, CREB activation via Ca^2+^-dependent CaMKII might also occur downstream of Piezo1 channel activation.Nevertheless, since most of the CREB activation and IL-6 upregulation was inhibited by the inhibitor, it appears that CREB phosphorylation and IL-6 upregulation in Piezo1-mediated Ca^2+^ signaling in HConjECs are predominantly attributable to p38 MAPK-dependent signaling. Importantly, the promoter of the human *IL-6* gene contains a cAMP-response element (CRE) where activated CREB specifically binds to regulate transcription. This supports our findings that the upregulation of IL-6 expression in HConjECs via Piezo1 channel activation was mediated through the p38 MAPK-CREB pathway [40].

We also confirmed the activation of p38 MAPK and CREB, along with an upregulation of IL-6 expression, following Piezo1 activation *in vivo*.These events occurred more rapidly than *in vitro*, necessitating shorter sampling intervals after Yoda1 eye drop administration. The findings are still preliminary, but MPO-positive neutrophil infiltration into the rat conjunctival epithelium was observed 24 hours post-Piezo1 activation. In contrast, the number of MBP-positive eosinophils in the conjunctival epithelium remained at basal levels at both 3 and 24 hours. Interestingly, Romac and colleagues have demonstrated that both applying pressure to the pancreatic duct and exposing it to Yoda1 induced pancreatitis in mice, accompanied by increased MPO concentrations in the tissue, supporting the findings of our study [41].

These granulocytes are crucial for the ocular surface’s defense as it interfaces with the external environment, acting as a primary barrier against external substances and pathogens. Neutrophils are primarily responsible for rapid responses to bacterial and fungal infections, whereas eosinophils play significant roles in allergic reactions and parasitic infections. With the experiments using various allergens and infectious agents, therefore, mechanical stress-induced Piezo1 activation in the conjunctival epithelium may trigger an inflammatory response similar to that seen during bacterial or fungal infections.

Although neutrophils express IL-6 receptors, it remains controversial whether IL-6 functions as a chemoattractant [42]. Given that IL-8 is known to induce neutrophil infiltration into tissues [43,44], Piezo1 activation-induced IL-8 production might have facilitated neutrophil infiltration into the conjunctival epithelium. However, tracking IL-8 protein expression in HConjECs was challenging in this study. Further studies are necessary to explore the connections between cytokine production in conjunctival epithelial cells, triggered by mechanical stress through Piezo1 activation, and neutrophil infiltration into the conjunctival epithelium.

As mentioned above, there is direct evidence supporting the induction of inflammatory cytokine expression due to eye-rubbing in humans.Balasubramanian *et al*. have reported significant increases in the levels of pro-inflammatory cytokines in the tear fluid following eye-rubbing. Their findings demonstrate elevated levels of IL-6 (1.24 ± 0.98 vs. 2.02 ± 1.52 pg/ml) and TNF-α (1.16 ± 0.74 vs. 1.44 ± 0.66 pg/ml), as well as MMP-13 (51.9 ± 34.3 vs. 63 ± 36.8 pg/ml, p = 0.006) [19]. Moreover, many studies related to keratoconus which is associated with eye rubbing, and the use of contact lenses—both of which are considered to induce mechanical stress on the ocular surface tissues—commonly report increased concentrations of IL-6, IL-1β, TNF-α, and MMP-9 in tear fluid compared to healthy subjects [[14], [15], [16], [17]]. These findings, together with the results of the current study, strongly suggest that mechanical stimulation of ocular surface tissues induces a Piezo1-dependent increase in inflammatory cytokines in the conjunctival epithelium, making it a significant risk factor for ocular inflammation.

## 5 Conclusion

Nearly everyone knows that we should not rub our eyes, often attributing this advice to the risk of infection from unclean fingers or potential damage to the ocular surface. However, our study demonstrated that eye-rubbing can induce the production of inflammatory cytokines in the conjunctiva, leading to neutrophil infiltration. Our result of neutrophil infiltration and the result that cellular responses observed after Piezo1 activation were similar to those triggered by fungal exposure, suggesting that mechanical stimulation of the conjunctiva can elicit an inflammatory response akin to an infection. This implies that ocular itching caused by infections may prompt eye-rubbing, thereby enhancing the inflammatory defense mechanisms. However, our findings with other studies indicate that habitual eye-rubbing, or excessive eye-rubbing induced by allergic symptoms, can result in an overactive inflammatory response, increasing the risk of developing or exacerbating ocular inflammatory or inflammation-related diseases such as keratoconus. Thus, raising awareness about the risks of persistent eye-rubbing is crucial for preventing the onset and progression of ocular inflammation.

## Declaration of competing interest

The authors have no conflicts of interest to disclose.

## Acknowledgement

We would like to express our sincere gratitude to Dr. Yasunori Takayama of Showa University for his invaluable advice and guidance on the experiments related to Piezo1. His insights and expertise were instrumental in the success of this research.

## Notes

### Competing Interest Statement

The authors have declared no competing interest.

